# Nucleic Acid Cleavage with a Hyperthermophilic Cas9 from an Unculturable Ignavibacterium

**DOI:** 10.1101/555169

**Authors:** Stephanie Tzouanas Schmidt, Feiqiao Brian Yu, Paul C. Blainey, Andrew P. May, Stephen R. Quake

## Abstract

CRISPR-Cas9 systems have been effectively harnessed to engineer the genomes of organisms from across the tree of life. Nearly all currently characterized Cas9 proteins are derived from mesophilic bacteria, and canonical Cas9 systems are challenged by applications requiring enhanced stability or elevated temperatures. We discovered IgnaviCas9, a Cas9 protein from a hyperthermophilic Ignavibacterium identified through mini-metagenomic sequencing of samples from a hot spring. IgnaviCas9 is active at temperatures up to 100 °C *in vitro*, which enables DNA cleavage beyond the 44 °C limit of *Streptococcus pyogenes* Cas9 (SpyCas9) and the 70 °C limit of both *Geobacillus stearothermophilus* Cas9 (GeoCas9) and *Geobacillus thermodenitrificans* T12 Cas9 (ThermoCas9). As a potential application of this enzyme, we demonstrated that IgnaviCas9 can be used in bacterial RNA-seq library preparation to remove unwanted cDNA from 16s ribosomal rRNA (rRNA) without increasing the number of steps, thus underscoring the benefits provided by its exceptional thermostability in improving molecular biology and genomic workflows. Taken together, IgnaviCas9 is an exciting addition to the CRISPR-Cas9 toolbox and expands its temperature range.

## Introduction

The application of clustered regularly interspaced short palindromic repeats (CRISPR) and CRISPR-associated (Cas) proteins has revolutionized molecular biology by making genome editing facile in both prokaryotes and eukaryotes (Jinek, Cong). Constituting the heritable and adaptive immune system of prokaryotes, CRISPR-Cas9 systems are present in archaea and bacteria from diverse environments (Koonin). A wide variety of CRISPR-Cas9 systems exist, and Class 2 systems, particularly type-II systems, have been well characterized and broadly implemented in part because these systems rely on a single effector protein, Cas9, and an RNA duplex, which can be replaced by a single guide RNA (sgRNA). CRISPR-Cas9 systems, particularly that from *Streptococcus pyogenes*, have been leveraged to edit genomes across organisms, create new tools for sequencing applications, and more (Wang).

Nearly all Cas9 proteins have been derived from mesophilic hosts, and thus cannot be used in applications in which elevated temperatures and robust stability are required. Recently, two Cas9 proteins from thermophiles were reported, providing enhanced stability in *in vivo* environments and enabling genome editing in thermophilic organisms (Harrington, Mougiakos). These two proteins, GeoCas9 and ThermoCas9, were identified by sequencing environmental samples, and their hosts live at temperatures of 65 and 70 °C, respectively.

More broadly, thermostable enzymes have led to important advances in biotechnology such as the polymerase chain reaction (PCR). The ability to use CRISPR-Cas9 at even higher temperatures than those reported would enable exciting new molecular biology techniques. Our group recently used mini-metagenomic sequencing to discover a distinct and novel CRISPR-Cas9 system in a hyperthermophilic bacterium from Yellowstone National Park’s Lower Geyser Basin whose temperatures exceed 90 °C (Yu).

Motivated by the potential of CRISPR-Cas9 at such elevated temperatures, we expressed, purified, and characterized this type II-C Cas9 protein, which we call IgnaviCas9. IgnaviCas9 is active at temperatures up to 100 °C; the active temperature range of IgnaviCas9 is the largest yet reported for a CRISPR-Cas9 system, opening myriad molecular biology applications. As one potential application, we demonstrate the reduction of undesirable 16s rRNA library molecules in bacterial samples being prepared for RNA-Seq.

## Results

### Identification, phylogenetic characterization, and expression of IgnaviCas9

Microfluidic mini-metagenomic sequencing of a sediment sample from the Lower Geyser Basin of Yellowstone National Park yielded a complete metagenome assembled genome. Comprised of a single 3.4 Mb contig representing a novel lineage in the *Ignavibacteriae* phylum, this genome contained a full CRISPR array. The temperature of the sample was recorded as 55 °C and that of the hot spring as >90 °C. That genome editing could take place in a bacterium residing in a near-boiling ecosystem intrigued us, and we examined the sequence of the CRISPR array for insights into its properties.

The CRISPR array contained a Cas9 protein, Cas1 protein, and Cas2 protein along with 38 unique spacers. The absence of a Csn2 and Cas4 protein suggested that the Ignavibacterium possessed a type II-C system (Mir), which was confirmed by phylogenetic comparison of IgnaviCas9 to other type II Cas9 proteins (Fig. 1A). Briefly, multiple sequence alignment of amino acid sequences of representative type II Cas9 proteins was performed using MAFFT, and a maximum-likelihood phylogenetic tree was constructed using RAxML with the PROTGAMMALG substitution model and 100 bootstrap samplings (Burstein). Like GeoCas9 and ThermoCas9, IgnaviCas9 is a type II-C Cas9. However, IgnaviCas9 is located in an entirely different clade, suggesting that IgnaviCas9 is highly divergent from the two thermostable Cas9s reported thus far. The *in vitro* validated type II-C Cas9 to which it is most similar is that of *Parvibaculum lavamentivorans* (Ran), a mesophilic bacterium with an optimal growth temperature of 30 °C (Schleheck).

**Figure 1.**
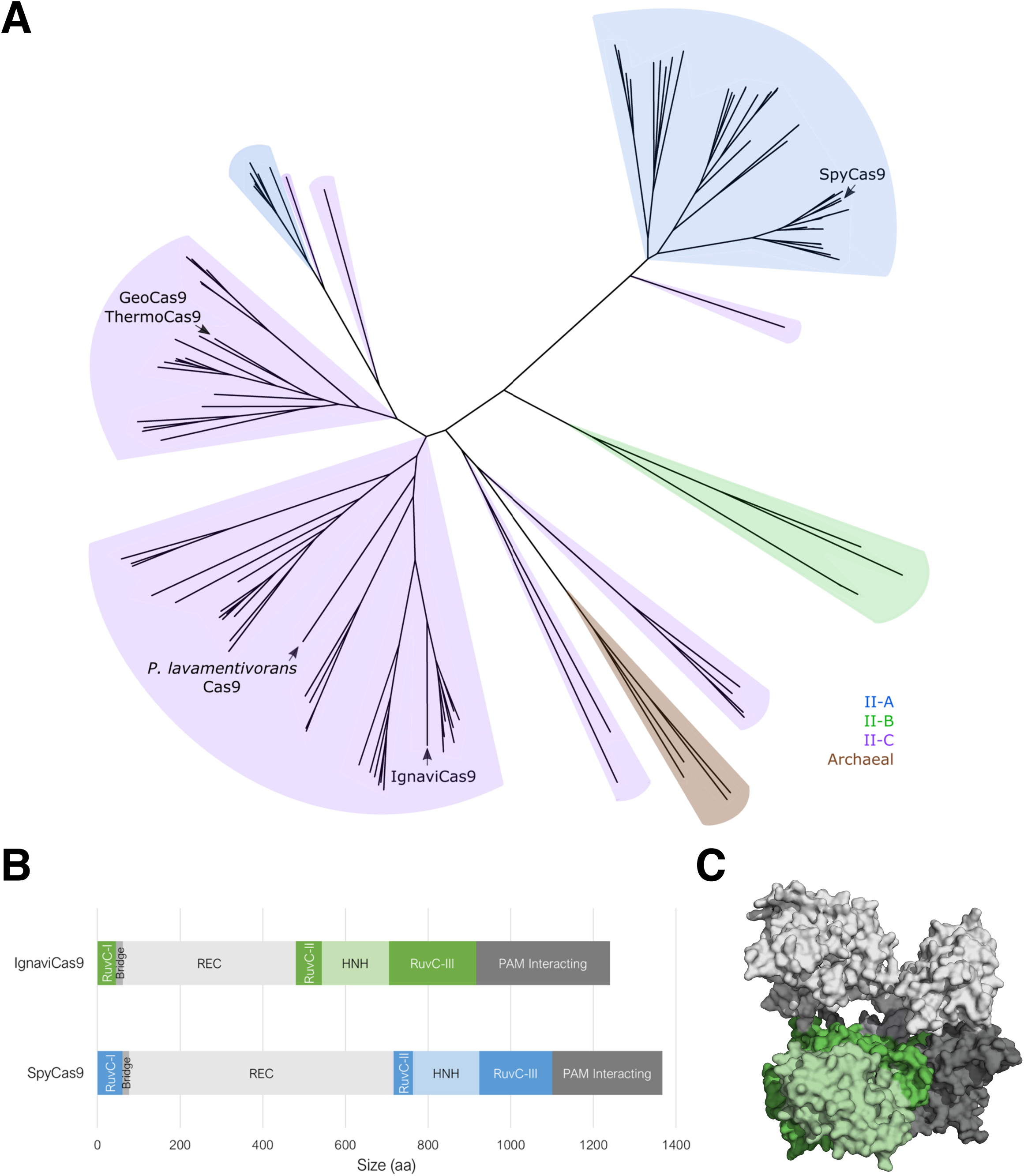
Phylogenetic classification and structural prediction of IgnaviCas9. A) Phylogenetic tree of representative Cas9s from Type II systems. B) Architectural domains of IgnaviCas9 and SpyCas9 where REC is the recognition lobe. C) A homology model of IgnaviCas9 with colors indicating the domains annotated in 1B. The model was generated using using Phyre2 (Kelley).

At 1241 amino acids long, IgnaviCas9 is certainly shorter than SpyCas9 (1368 amino acids) but longer than ThermoCas9 (1082 amino acids) or GeoCas9 (1087 amino acids). Through homology modeling and sequence alignment, the smaller size of IgnaviCas9 compared to SpyCas9 was found to arise from its reduced REC lobe (Fig. 1B), which is consistent with other smaller Cas9s (Ran). While larger than other *in vitro* validated type II-C Cas9 proteins, IgnaviCas9 is shorter than SpyCas9, which can be advantageous for applications involving its delivery via adeno-associated viruses (Wu).

Having examined IgnaviCas9 phylogenetically, we sought to produce and test the protein. Because the bacterium IgnaviCas9 comes from has not been isolated and is likely unable to be grown in standard lab culture conditions, we codon-optimized the sequence of IgnaviCas9 and cloned it into a Cas9-expression vector. BL21 *E. coli* cells were transformed with this plasmid and cultured to express IgnaviCas9. Subsequent purification provided 12 mg of IgnaviCas9 from 4 L of culture for downstream experiments.

### IgnaviCas9 sgRNA engineering

That IgnaviCas9 falls within the type II-C classification proved helpful in designing its sgRNA based on computational prediction of its crRNA and tracrRNA from the CRISPR array sequence. This approach was necessary, since the bacterium from which IgnaviCas9 was isolated not available and unlikely to be culturable. We looked for combinations of potential crRNA and tracrRNA sequences that together allowed for the formation of the lower stem, bulge, upper stem, nexus, and hairpin features (Fig. 2A).

**Figure 2.**
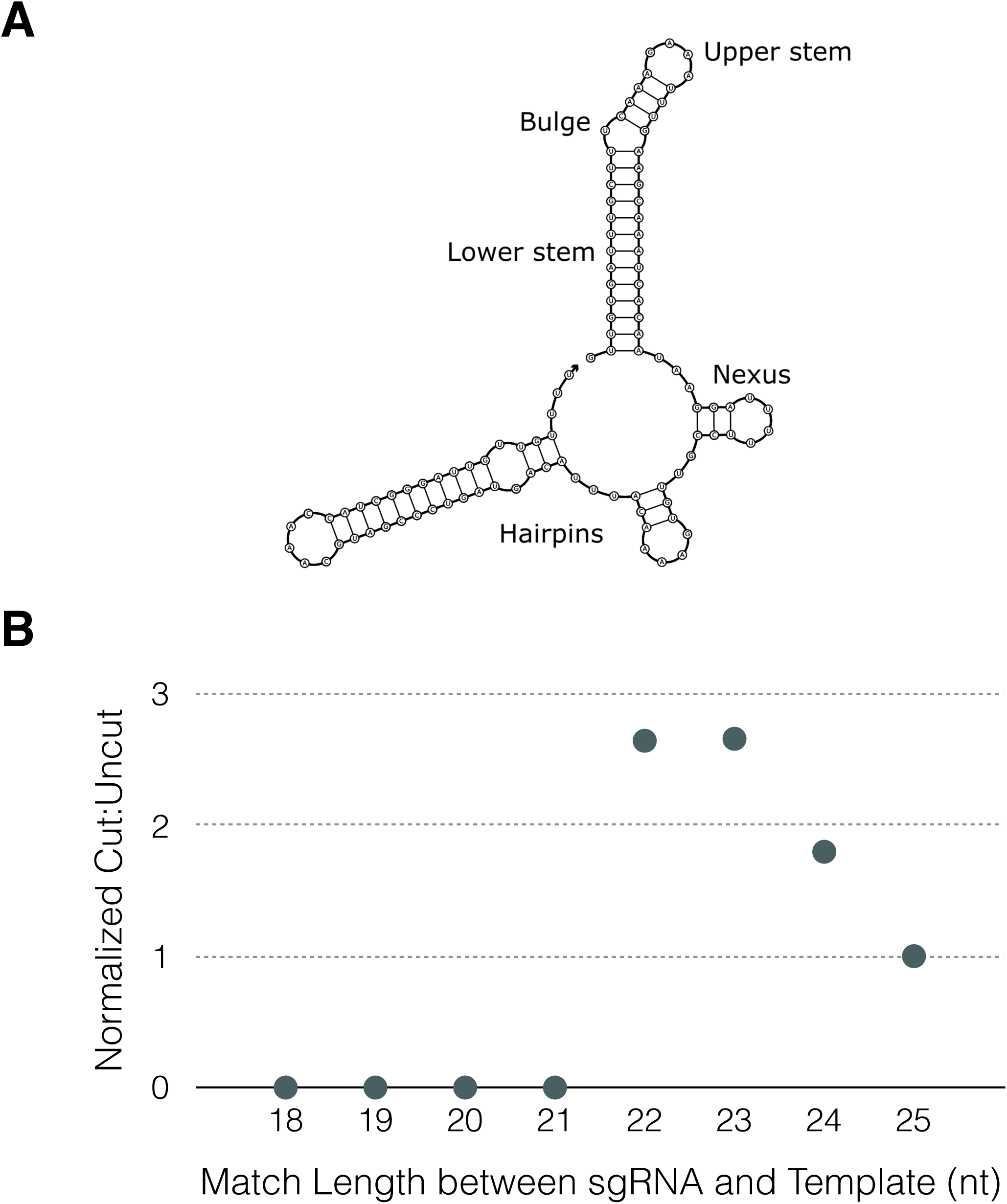
sgRNA structure and spacer length. A) Representation of the determined sgRNA with important structural features labeled. B) Testing of the preferred spacer length was conducted by comparing cleavage at 52 °C of templates targeted by truncated versions of the initial spacer. The cut-to-uncut-ratio was normalized to that corresponding to 25 nt (length used for preliminary experiments).

RNA secondary structure prediction of the designed sgRNA showed that all desired features remained present at temperatures of 60 °C for default NUPACK program settings, underscoring the potential of IgnaviCas9 to cleave DNA at temperatures outside of the mesophilic range. We transcribed the sgRNA sequence preceded by 25 nt of spacer sequence for use in preliminary experiments.

### IgnaviCas9 PAM determination and sgRNA-spacer match length refinement

The protospacer adjacent motif (PAM), the sequence directly downstream of a nucleic acid target cleavable by CRISPR systems, varies between different species and prevents the host genome from being attacked (Mojica). As an initial approach, we designed double-stranded linear DNA containing a spacer sequence followed by a PAM from an *in vitro* validated type II-C CRISPR system (Mir). We performed cleavage assays by incubating the assorted DNA substrates with a ribonucleoprotein complex (RNP) of IgnaviCas9 and sgRNA targeting the spacer sequence at 52 °C for 30 minutes. We found that IgnaviCas9 cleaved the DNA substrate with the PAM CCACATCGAA, containing the NNNCAT motif from *P. lavamentivorans* (Fig. 3A). A control reaction was used as a point of reference and differed in that the sgRNA included contained a scrambled version of the spacer sequence.

**Figure 3.**
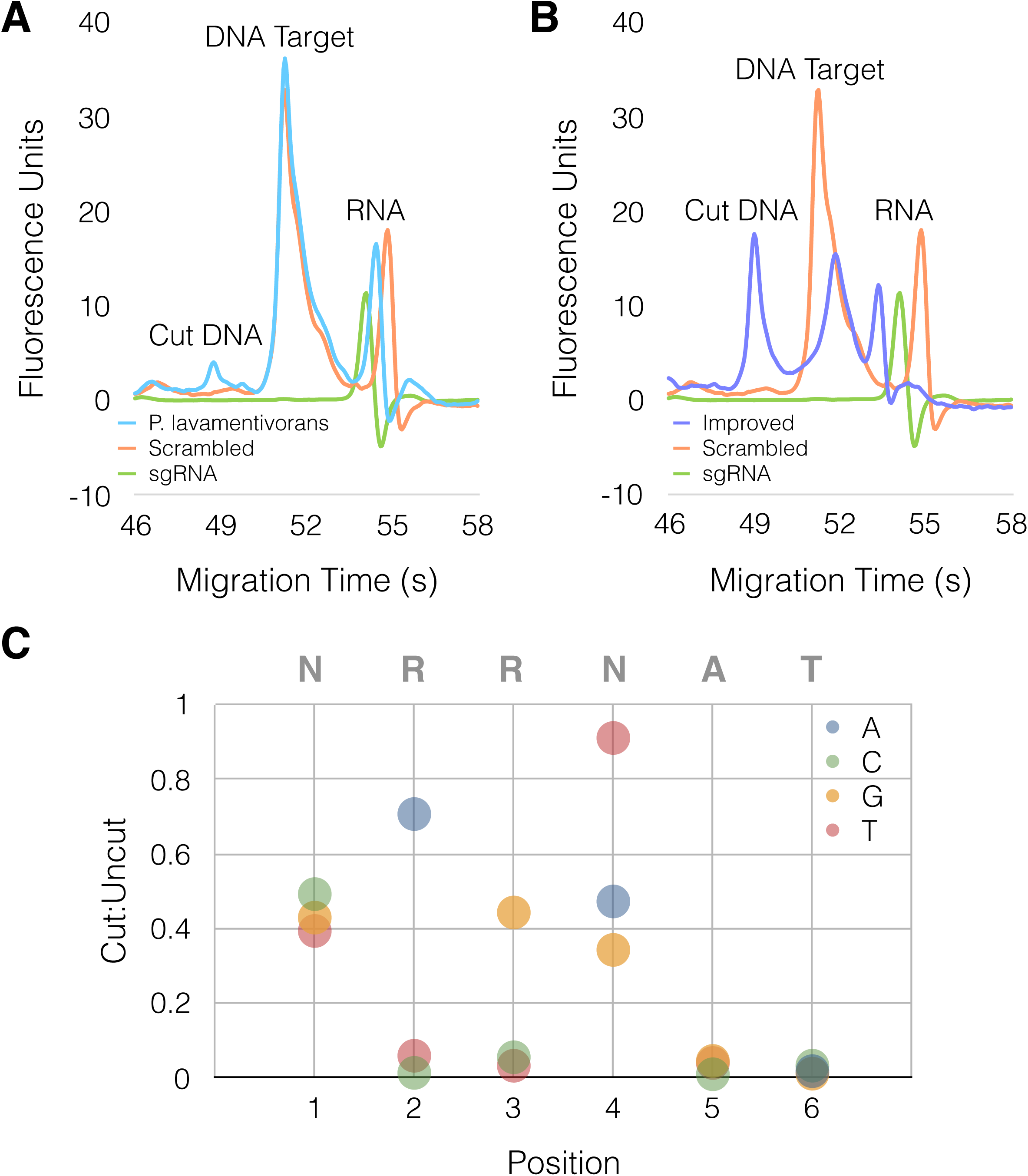
PAM determination. A) Electropherogram showing cleavage of template containing the PAM from *P. lavamentivorans* compared to control reaction with scrambled sgRNA and to sgRNA from experimental condition. B) Electropherogram showing cleavage of template containing the PAM from *P. lavamentivorans* with adjustments informed by leads from bulk sequencing data. Curves from control reaction with scrambled sgRNA and from experimental condition sgRNA are included for comparison. C) Performance of IgnaviCas9 in cleaving DNA templates with the indicated substitutions at the specified positions for the starting sequence of AGACAT. Substitutions abolishing cleavage activity enabled PAM refinement.

We then used the 38 spacers found in the IgnaviCas9 CRISPR array to isolate possible protospacers from the environmental sample in which IgnaviCas9 was found. By using BLAST to search the environmental sequences, we collected 10 bp sequences flanking the spacer that were different from the repeat sequence by an edit distance of at least 5. The sequence logo created using unique sequences meeting these criteria suggested that the PAM was likely to be adenine-rich (Fig. S1); positions 7 through 10 are not shown as they were ultimately determined to not impact the PAM. We designed a new DNA substrate by modifying the aforementioned DNA substrate that was cut by IgnaviCas9 to include AGACATGAAA, an adenine-rich version of the *P. lavamentivorans* PAM. In a cleavage reaction performed as before, IgnaviCas9 was able to better cleave the DNA substrate containing the refined PAM (Fig. 3B).

We finalized the PAM recognized by IgnaviCas9 by testing DNA substrates containing the adenine-rich *P. lavamentivorans* PAM with single nucleotide substitutions at each of the 10 positions directly downstream of the spacer. To concisely convey the performance of IgnaviCas9 in cleaving the DNA template, we calculated a metric we term the cut-to-uncut ratio by dividing the area under the larger cut peak by the area under the uncut peak (Fig. S2). Disruption of IgnaviCas9 cleavage by a particular substitution demonstrated that the position of the substitution was important to the PAM and that the nucleotide was not part of the PAM. We found that NRRNAT is the PAM recognized by IgnaviCas9; all substitutions at positions past the sixth bp downstream of the spacer sequence were tolerated (Fig. 3C).

Having established IgnaviCas9’s PAM, we varied the length of spacer included in the sgRNA to determine which lengths were optimal. We demonstrated that IgnaviCas9 cleaves DNA when the sgRNA includes spacer lengths of 22 to 25 nt, with improved performance for 22 or 23 nt spacer lengths (Fig. 2B). Cleavage does not occur for sgRNA with shorter spacer lengths. The spacer sizes IgnaviCas9 prefers overlap with those favored by ThermoCas9 (19 to 25 nt) and GeoCas9 (21 or 22 nt) but are slightly larger than the 20 nt spacer length typically used with SpyCas9.

### Active temperature range assessment

By conducting the PAM determination experiments at 52 °C, we confirmed that IgnaviCas9 is active at temperatures above those of the active range of SpyCas9, which has been reported as between 20 and 44 °C (Wiktor, Mougiakos). To quantify the efficiency of IgnaviCas9 in cleaving DNA, we defined its efficiency as the area under the larger cut peak from the test condition divided by the area under the uncut peak from the scrambled control (Fig. S3). We characterized the temperature range over which IgnaviCas9 is active by performing cleavage assays between 31 and 100 °C (Fig. 4A). We found that its performance in cutting various DNA targets, including longer templates like plasmid DNA, extended across the range tested, which reaches beyond the upper active temperature limit of other thermostable Cas9 proteins. That IgnaviCas9 remains active at high temperatures and across a wide thermal range (Fig. 4B) indicates that it is particularly stable and suggests that it is likely more specific in its targeting than SpyCas9. This is consistent with the lower mismatch tolerance of other thermostable Cas9 proteins compared to SpyCas9 (Harrington, Mougiakos).

**Figure 4.**
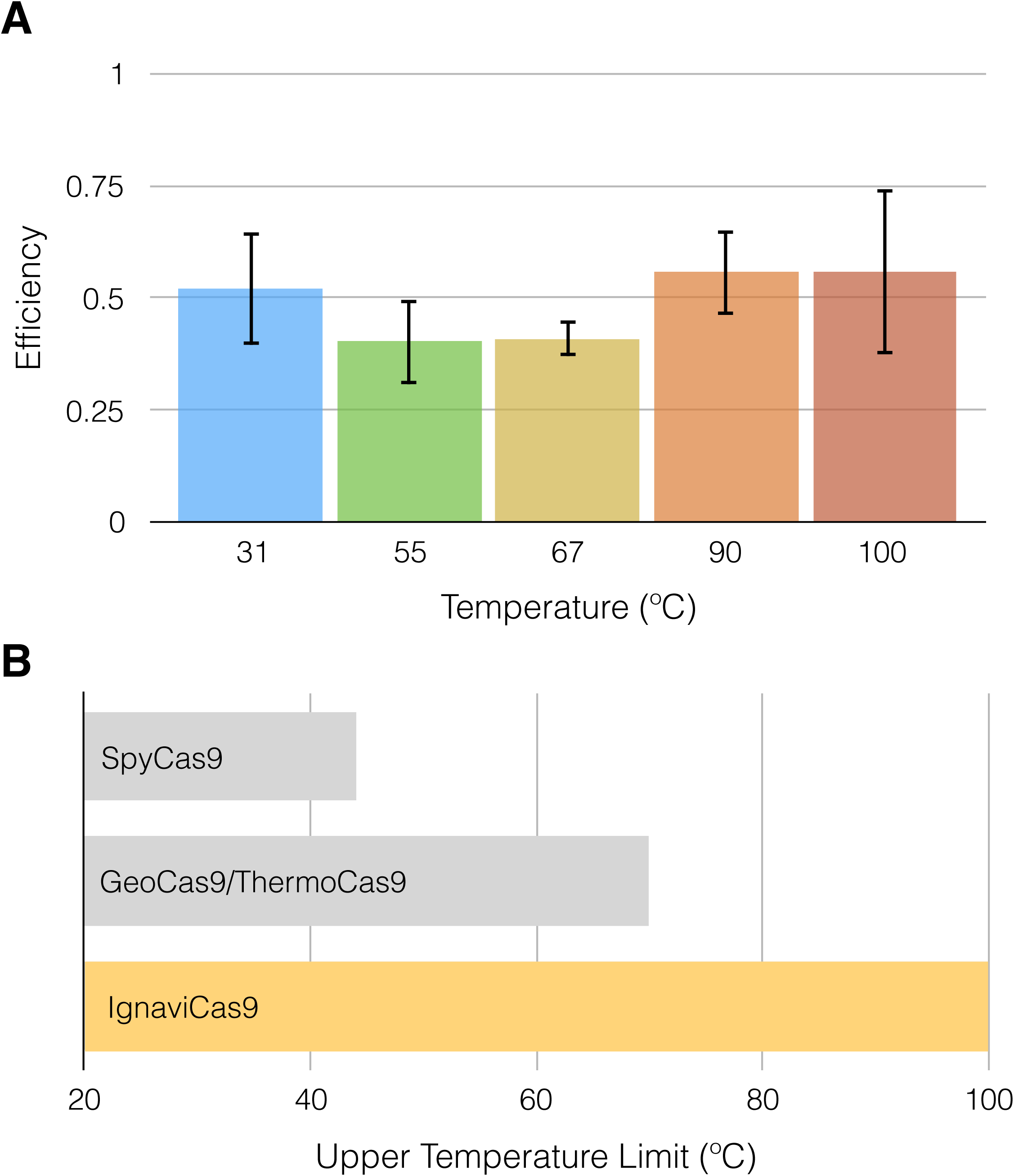
Temperature evaluation of IgnaviCas9. A) The efficiency of IgnaviCas9 in cleaving DNA templates is compared over a range of temperatures. The average and standard deviation at each temperature tested is shown (n=3). B) The upper temperature limit of Cas9 homologs.

### Implementation of IgnaviCas9 to remove undesired amplicons

The wide active temperature range of IgnaviCas9 is a unique property that can be harnessed for a host of molecular biology applications. In particular, its activity at both moderate and high temperatures led us to consider how it could be integrated into molecular biology and genomic workflows to eliminate undesired amplicons. Given our prior work in mini-metagenomics, we were interested in leveraging IgnaviCas9 to reduce the amplification of library molecules derived from 16s rRNA in bacterial RNA-Seq.

When performing RNA-seq of actively growing bacterial strains or generating metatranscriptomic data from environmental samples, reads from 16s rRNA genes are typically highly abundant and reduce sequencing bandwidth of expression profiles of interest. With this in mind, we deployed IgnaviCas9 during the PCR step of the sequencing library preparation workflow to cleave library fragments derived from 16s rRNA, thus reducing their presence in the final library without adding steps to the workflow. While a mesophilic Cas9 could be used in an additional workflow step prior to amplification to cleave undesired fragments in a library, concurrent amplification and targeted depletion offers multiple opportunities to cleave the target and prevent its exponential amplification.

To this end, we designed sgRNA to target cDNA resulting from 16s rRNA and included IgnaviCas9 complexed to these sgRNAs in a PCR. Through sequencing, we demonstrated that, compared to the starting library, IgnaviCas9 targeting during amplification reduced the contribution of libraries derived from 16s rRNA, thus enriching the portion containing transcripts of interest (Fig. 5). More broadly, our approach could be used to eliminate other unwanted amplicons, e.g. primer dimers, as they are generated. Such implementations of IgnaviCas9 underscore its utility in improving widely used existing techniques in genomics and molecular biology.

**Figure 5.**
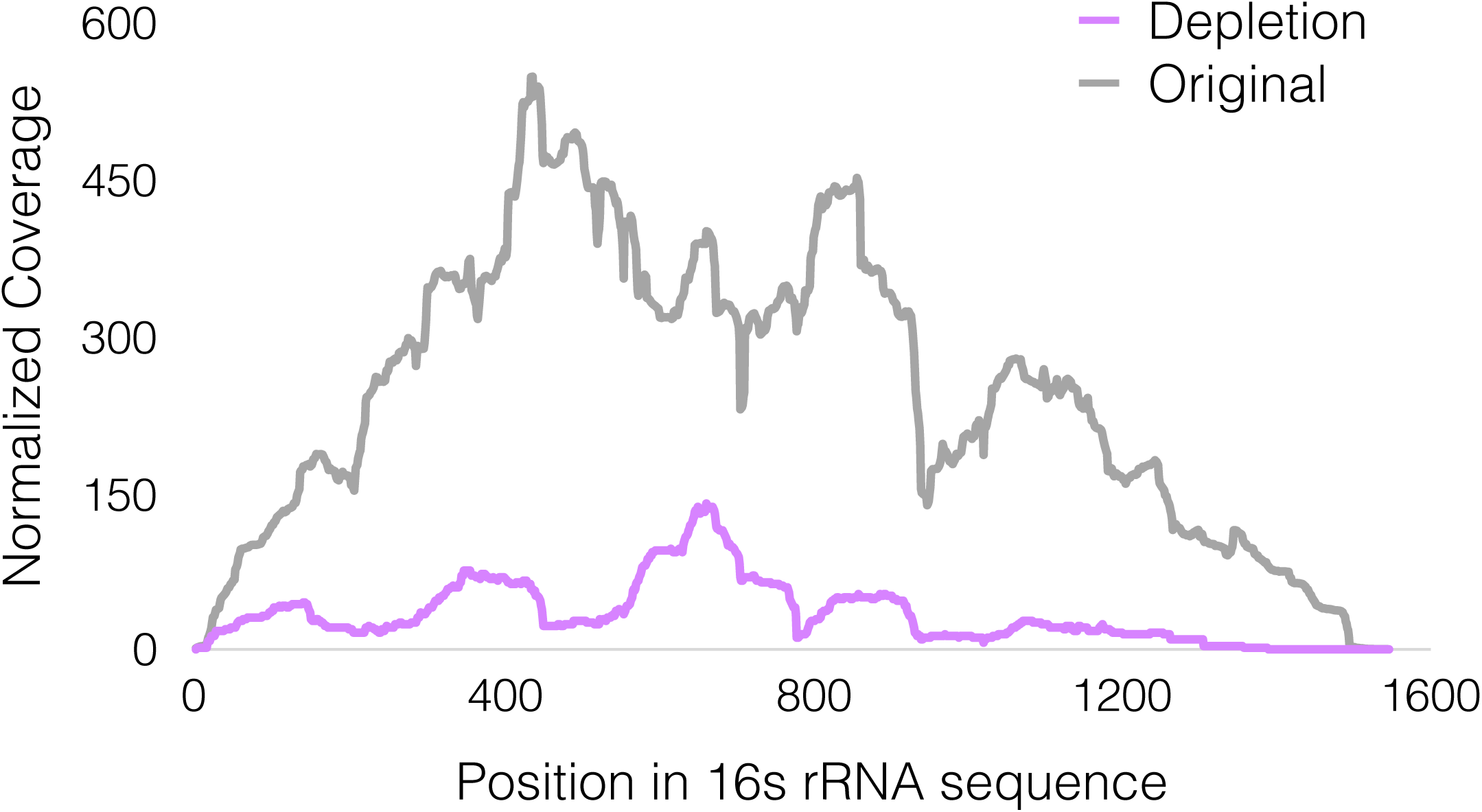
Reduction of targeted sequence by IgnaviCas9. Coverage plot for 16s rRNA sequence targeted by IgnaviCas9 during PCR amplification. Normalized coverage given as per-base coverage divided by average whole genome coverage.

## Discussion

Taken together, our work expands the possible range of CRISPR-Cas9 to temperatures as high as 100 °C. While nearly all characterized Cas proteins are active at mesophilic temperatures, IgnaviCas9 is clearly hyperthermophilic while also possessing activity at lower temperatures as well. Its natural propensity to cleave DNA across such a wide temperature range circumvents the need for protein engineering to create such a Cas9 and underscores the value of environmental metagenomics. By exploring the rich diversity of microbes present in extreme environments were we able to identify a promising CRISPR-Cas9 system for further study and use. IgnaviCas9 is highly thermostable, which enables a wide variety of important applications. The highest previously reported active temperature for a Cas9 is 70 °C.

That IgnaviCas9 is able to bind and cleave DNA at such high temperatures underscores its stability, a feature that could make it well suited for *in vivo* use. In particular, increased stability suggests that IgnaviCas9 will have a longer lifetime in plasma than those of canonical variants and thus, will be more effective for applications like gene therapies (Long) or lineage tracing in complex organisms (Schmidt). While organisms dwelling at higher temperatures are typically unicellular microorganisms, these microbes can catalyze important high-temperature industrial processes like fermentation. The improved ability to engineer thermophilic bacteria by means of IgnaviCas9 will facilitate the development and broader implementation of these processes. Finally, the ability to edit DNA at elevated temperatures beyond the active range of previously reported Cas9s is key to both improving existing and creating new molecular biology applications in which temperature serves as a means of control.

## Methods

### IgnaviCas9 identification, expression, and purification

IgnaviCas9 was found through mini-metagenomic sequencing of a sediment sample taken from Mound Spring in the Lower Geyser Basin area of Yellowstone National Park under permit YELL-2009-SCI-5788. The sample was placed in 50% ethanol in a 2 mL tube without any filtering and kept frozen until returning from Yellowstone to Stanford University, at which time tubes containing the samples were transferred to −80°C for long term storage.

To compare IgnaviCas9 to other Cas9s (Burnstein), multiple sequence alignment of type II Cas9s was performed using MAFFT (Katoh), and a maximum-likelihood phylogenetic tree was constructed using RAxML with the PROTGAMMALG substitution model and 100 bootstrap samplings (Stamatakis).

Its DNA sequence was codon-optimized for expression in *E. coli* and then synthesized (Integrated DNA Technologies). The resulting DNA was cloned into a pET-based vector with an N-terminal hexahistidine, maltose binding protein, and tobacco etch virus sequence and C-terminal nuclear localization sequences.

IgnaviCas9 was expressed in BL21 strain *E. coli* (Agilent). After cultures reached an OD_600 nm_ of 0.5, expression was induced by adding IPTG to give a final concentration of 0.5 mM. The cultures were allowed to incubate for 7 hours at 16 °C. Cells were harvested via centrifugation, and IgnaviCas9 was purified using ion exchange and size exclusion chromatography per previously described methods (Gu). IgnaviCas9-containing fractions were pooled, supplemented with glycerol to a final concentration of 50%, and stored at −80 °C until used.

### sgRNA design and transcription

The crRNA and tracrRNA were identified from the IgnaviCas9 CRISPR locus by searching for complementarity between candidate sequences that allowed for the formation of the requisite features when linked by a 5’-GAAA-3’ tetraloop (Briner). Possible sgRNA sequences were tested through secondary structure prediction using NUPACK (Zadeh).

DNA corresponding to the sgRNA including the target of interest was placed under control of a T7 promoter and synthesized (Integrated DNA Technologies). sgRNAs were transcribed using the MEGAshortScript T7 Transcription Kit (Thermo Fisher Scientific) with overnight incubation and purified using the MEGAclear Transcription Clean-Up Kit (Thermo Fisher Scientific).

### *In vitro* cleavage assays

The purified IgnaviCas9 and transcribed sgRNA were used to cleave DNA targets at desired temperatures. Templates approximately 100 bp long used in the PAM determination experiments and temperature range testing were synthesized (Integrated DNA Technologies). Plasmid templates for additional temperature range testing were generated by linearizing the pwtCas9 plasmid (Qi) using XhoI (New England Biolabs).

IgnaviCas9 and the appropriate sgRNA were incubated together in reaction buffer at 37 °C for 10 minutes before adding the DNA target to the reaction. The reaction was then incubated at the specified temperature for 30 minutes. The final composition of each reaction was 5 nM substrate DNA, 100 nM IgnaviCas9, 150 nM sgRNA, 20 mM Tris-HCl pH 7.6, 100 mM KCl, 5 mM MgCl_2_, 1mM DTT, and 5% glycerol (volume per volume).

Each reaction was quenched using 6x Quench Buffer (15% glycerol, 100 mM EDTA) and then underwent Proteinase K digestion at room temperature for 20 minutes before being loaded into a chip for fragment analysis using the Bioanalyzer (Agilent).

### 16s rRNA depletion in bacterial RNA-Seq libraries

Four different sgRNAs were designed to target cDNA arising from 16s rRNA sequences. The sgRNA complexed with IgnaviCas9 as described above was added to cDNA derived from *E. coli* RNA that underwent reverse transcription and amplification using the ScriptSeq Complete Gold Kit for Epidemiology (Epicentre).

The HiFi HotStart ReadyMixPCR Mix (KAPA) was used for the combined amplification and targeted depletion reaction, comprised of 25 μL HiFi HotStart ReadyMixPCR Mix, 1 μL ScriptSeq Index PCR Primer (Epicentre), 1 μL Reverse PCR Primer (Epicentre), 1 ng of cDNA library, 2.5 μL of 5.5 μM IgnaviCas9, 15 μL of 1400 nM sgRNA, 5 μL of IgnaviCas9 reaction buffer, and water to a total volume of 50 μL. The cycling protocol used was as follows: 95 °C for 3 minutes, 30 cycles of 98 °C for 20 seconds and 75 °C for 30 seconds, and 72 °C for 1 minute.

A MiSeq Micro run was performed to sequence the original library and the test reaction that underwent concurrent amplification and targeted depletion. Resulting sequence reads were quality-filtered and trimmed using bbduk, aligned to the 16s rRNA sequence using bowtie2, and then sorted and indexed using samtools. Positional sequence coverage was determined using bedtools and subsequently compared between samples by normalizing to the average whole genome coverage in each sample.

## Additional Information

### Acknowledgements

We thank Emily Crawford (Chan Zuckerberg Biohub) for the plasmid used in protein expression and Richard Pfuetzner (Stanford University) for assistance with protein purification. We would also like to thank Anastasia Nedderton (Stanford University) and National Park Service staff at Yellowstone National Park including Research Coordinator Christie Hendrix for assistance with different aspects of the sample collection, preservation, and characterization processes.

S.T.S. is supported by the Fannie and John Hertz Foundation Fellowship, National Science Foundation Graduate Research Fellowship, Siebel Scholarship, and Gabilan Stanford Graduate Fellowship. P.C.B was supported by the Burroughs Welcome Fund via a Career Award at the Scientific Interface. This work was supported by a Boundaries of Life program grant from the Templeton Foundation and by the Chan Zuckerberg Biohub (S.T.S., B.Y., S.R.Q.).

### Author contributions

Conceived and designed the experiments: STS FY APM SRQ. Contributed reagents/materials/analysis tools: PCB SRQ. Performed the experiments: STS FY. Analyzed the data: STS FY APM SRQ. Wrote the paper: STS SRQ.

## Supplementary Figure Captions

**Supplementary Figure 1.**
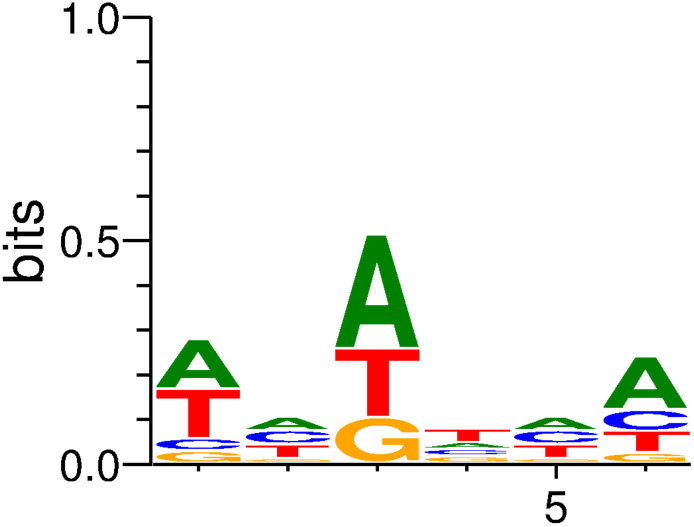
PAM informed by spacer search. Logo resulting from sequences flanking IgnaviCas9 CRISPR array spacers that were identified from bulk sequencing of environmental sample in which IgnaviCas9 was found.

**Supplementary Figure 2.**
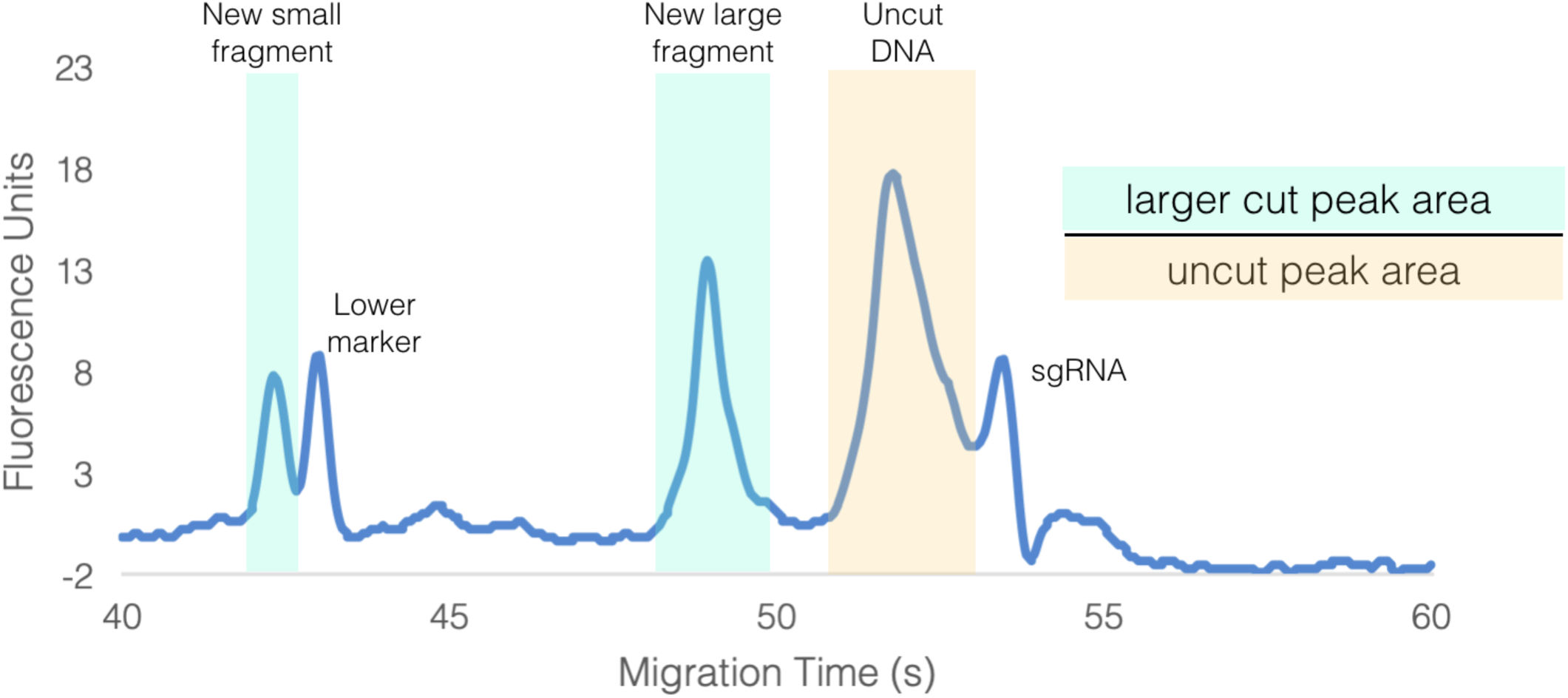
Cut:Uncut. Electropherogram showing notable peaks in cleavage reactions. The cut-to-uncut ratio is calculated by dividing the area under the larger cut peak by the area under the uncut peak.

**Supplementary Figure 3.**
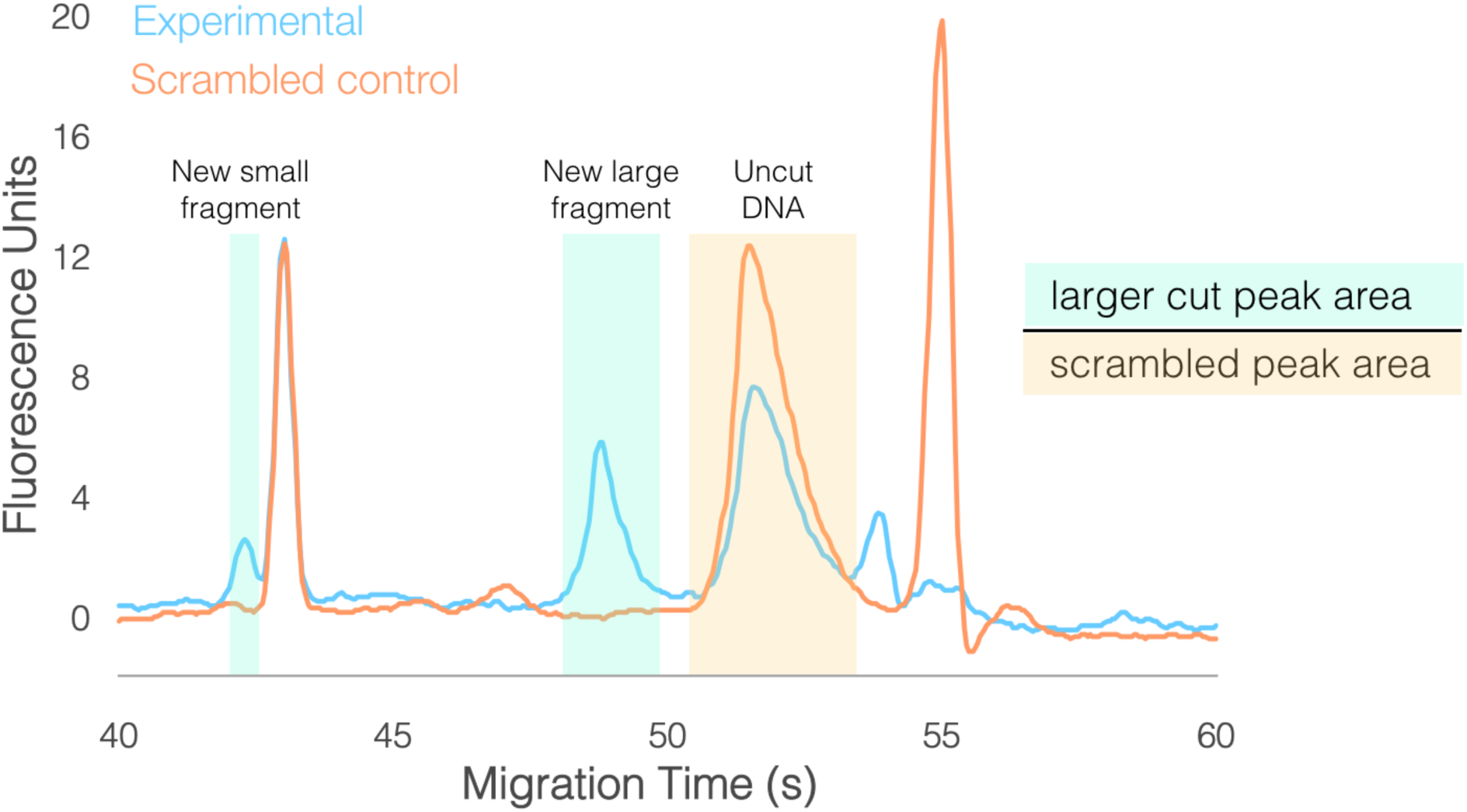
Efficiency. IgnaviCas9’s efficiency is calculated by dividing the area under the larger cut peak from the experimental reaction by the area under the uncut peak from the scrambled control.

